# Multiplexed Ion Beam Imaging Readout of Single-Cell Immunoblotting

**DOI:** 10.1101/2021.03.06.434187

**Authors:** Gabriela Lomeli, Marc Bosse, Sean C. Bendall, Michael Angelo, Amy E. Herr

## Abstract

Improvements in single-cell protein analysis are required to study the cell-to-cell variation inherent to diseases, including cancer. Single-cell immunoblotting (scIB) offers proteoform detection specificity, but often relies on fluorescence-based readout and is therefore limited in multiplexing capability. Among rising multiplexed imaging methods is multiplexed ion beam imaging by time of flight (MIBI-TOF), a mass spectrometry imaging technology. MIBI-TOF employs metal-tagged antibodies that do not suffer from spectra overlap to the same degree as fluorophore-tagged antibodies. We report for the first-time MIBI-TOF of single-cell immunoblotting (scIB-MIBI-TOF). The scIB assay subjects single-cell lysate to protein immunoblotting on a microscale device consisting of a 50- to 75-μm thick hydrated polyacrylamide (PA) gel matrix for protein immobilization prior to in-gel immunoprobing. We confirm antibody-protein binding in the PA gel with indirect fluorescence readout of metal-tagged antibodies. Since MIBI-TOF is a layer-by-layer imaging technique, and our protein target is immobilized within a 3D PA gel layer, we characterize the protein distribution throughout the PA gel depth by fluorescence confocal microscopy and find that the highest signal-to-noise ratio is achieved by imaging the entirety of the PA gel depth. Accordingly, we report the required MIBI-TOF ion dose strength needed to image varying PA gel depths. Lastly, by imaging ~42% of PA gel depth with MIBI-TOF, we detect two isoelectrically separated TurboGFP (tGFP) proteoforms from individual glioblastoma cells, demonstrating that highly multiplexed mass spectrometry-based readout is compatible with scIB.

---

Single-cell analysis tools tease apart the cell-to-cell variability driving many important biological processes, such as cancer and drug resistance^1^. Underlying cellular heterogeneity is differential protein expression in individual cells; this molecular heterogeneity includes differential proteoform expression. Proteoforms are highly similar – yet chemically distinct – proteins originating from a single gene. Proteoforms often have unique functions^2,3^. For instance, in breast cancer tumors expressing human epidermal growth factor receptor 2 (HER2), the presence of truncated HER2 proteoforms has been connected to a decrease in the effectiveness of antibody therapy^4^. Moreover, despite the importance of measuring multiple proteins from single cells (i.e., to interrogate molecular circuits^5^ or categorize cell types^6^), target multiplexing in single-cell protein assays remains a major challenge in analytical chemistry^5^.

Current single-cell proteomic tools lack the capability to provide both proteoform specificity and high target multiplexing. Identification of proteoforms with conventional immunoassays, such as immunohistochemistry (IHC) and flow cytometry, requires proteoform-specific antibodies. Given the diversity of proteoform species possible, proteoform-specific antibodies are sometimes unavailable^7^. Further, each new target-specific probe requires an additional detection channel in what is usually an already crowded multiplexed antibody panel. Moreover, immunoassay methods typically rely on fluorescence detection as a readout, which has limited multiplexing ability due to the spectra overlap of fluorophores^8^. Mass spectrometry has directly detected >1000 protein types with single-cell resolution, but existing single-cell mass spectrometry has low throughput^1^, which makes identification of rare cell types difficult. Moreover, current single-cell mass spectrometry utilizes a “bottom-up” approach, and proteoforms are often not distinguishable with bottom-up mass spectrometry due to measurement of peptides, not intact proteins^2^. Thus, an unmet need remains for a single-cell protein analysis tool that provides proteoform specificity and is amenable to multiplexed protein-target detection.

MIBI-TOF is a mass spectrometry imaging technique especially designed for multiplexing and has been used to simultaneously image dozens of protein targets from fixed tissue^9^. For MIBI-TOF, the sample of interest, typically a tissue slice, is first immunoprobed with metal-isotope-tagged antibodies with each metal-isotope providing a distinct detection channel for target multiplexing. Then, MIBI-TOF uses secondary ion mass spectrometry (SIMS) to rasterize the sample with a primary ion beam, which sputters elements from both the metal-tagged antibodies and the sample to a time-of-flight spectrometer, generating a high parameter image comprised of the mass spectrum of each pixel. Although powerful in terms of target multiplexing and single-cell resolution, MIBI-TOF of intact tissue slices requires proteoform-specific antibodies for proteoform detection.

scIB provides proteoform specificity by first separating proteins by size (single-cell western blot [scWB]^10^) or charge (single-cell isoelectric focusing [scIEF]^11^), which relaxes the requirement of proteoform-specific antibodies since proteoforms are spatially separated prior to immunoprobing. Multiplexing to 12 protein targets in single cells has been reported in scIB by using an antibody stripping and reprobing approach in which 1-3 protein targets are imaged at a time with immunofluorescence, followed by a chemical stripping step, and the strip and reprobe process is cyclically repeated for additional targets^12^. However, there is a ~75% drop in immunoprobed signal after just one round of stripping^13^. Such signal losses create a challenge to target multiplexing that requires multiple stripping rounds, especially for the detection of low abundance proteins. The primary mechanism of signal decrease during stripping and reprobing is loss of ~50% of immobilized protein during the first round of stripping^13^; therefore, it is of great interest to eliminate the need for stripping altogether to achieve higher multiplexing.

To this end, here we introduce scIB with a MIBI-TOF readout. We report the characterization and validation steps taken towards realizing this technology. Since metal-tagging increases antibody probe size, which could lead to increased size-exclusion from the PA hydrogel matrix, we verify that metal-tagged antibodies can bind to their target in a scIB assay. Then, we characterize the depth distribution of signal in the 3D hydrogel matrix used by scIB assays in order to determine the percentage of the sample depth that should be imaged with MIBI-TOF, since physical removal of the substrate is needed for detection. In SIMS, the number of sputtered ions, and therefore thickness of the sample that is rasterized away, is related to the ion dose delivered to the sample per unit area (referred to as ion dose, hereafter)^14^. Accordingly, we measure the gel depth rasterized with varying ion doses. Finally, we image isoelectrically focused tGFP proteoforms from single cells with both a fluorescence microarray scanner and MIBI-TOF, utilizing an ion dose that rasterized approximately 42% of the gel depth, to demonstrate that scIB assays are compatible with MIBI-TOF detection.

## EXPERIMENTAL SECTION

### Chemicals/Reagents

PA gels were cast on silicon wafers (WaferPro C04009) microfabricated with SU8 3050 photo-resist (MicroChem Y311075) using custom in-house-designed masks (CAD/ART Services) and coated with dichlorodimethylsilane (Sigma 440272). An Ultrapure Millipore filtration system provided deionized water (18.2 MΩ). 3- (trimethoxysilyl)propyl methacrylate (Sigma 440159), methanol (VWR BDH1135), and glacial acetic acid (Fisher Scientific A38S) were used for silanization of standard glass slides (VWR 48300-048). 30%T 29:1 acrylamide/bis-acrylamide solution (Sigma A3574), 1.5 M pH 8.8 TrisHCl (TekNova T1588), N-[3-[(3- benzoylphenyl)formamido]propyl] methacrylamide (BPMAC, custom synthesized by PharmAgra Laboratories), ammonium persulfate (APS, Sigma A3678), and N,N,N′,N′-tetra-methylethylenediamine (TEMED, Sigma T9281) were used for microwell PA gel polymerization used in both scWB and scIEF. scWB was conducted using sodium deoxycholate (Sigma D6750), sodium dodecyl sulfate (SDS, Sigma L3771), TritonX-100 detergent (Sigma X100), and premixed 10× Tris/glycine electrophoresis buffer (25 mM Tris, pH 8.3; 192 mM glycine, BioRad 1610734) for the cell lysis buffer. scIEF was conducted using the immobilines pKa 3.6 and pKa 9.3 acrylamido buffers (Sigma 01716, 01738), ZOOM^®^ Carrier Ampholytes pH 4-7 (Thermo Fischer Scientific ZM0022), 40%T 29:1 acrylamide/bis-acrylamide solution (Sigma A7802), urea (Sigma U5378), thiourea (Sigma T8656), 3- [(3- Cholamidopropyl)dimethylammonio]-1-propanesulfonate (CHAPS, Sigma RES1300C), digitonin (Sigma D141), UV photoinitiator 2,2-Azobis(2-methyl-N-(2-hydroxyethyl) propionamide) (VA086, Wako Chemicals 013-19342), an ABS electrophoresis device designed and printed in-house, graphite electrodes (Bio-Rad 1702980), GelSlick (Lonza 50640), borosilicate glass sheets (McMaster-Carr 8476K62), and 0.5 mm gel spacers (CBS Scientific MVS0510-R). The antibody probes used were primary rabbit-anti-tGFP antibody (Pierce PA5−22688,) and secondary polyclonal antibody AlexaFluor-647-labeled donkey-anti-rabbit (Invitrogen A-31573). Bovine serum albumin (BSA, A7030) was purchased from Sigma-Aldrich. Tris-buffered saline with Tween-20 (TBS-T, Cell Signaling Technologies 9997S) was used for gel incubation and wash steps.

### Antibody Conjugation

Holmium (Ho)-tagged (metal-tagged) primary rabbit-anti-tGFP antibody was prepared using the MIBItag Conjugation Kit (Ho) (Ionpath 600165), which includes diethylenetriaminepentaacetic acid (DTPA) polymer pre-loaded with Holmium for conjugation to the antibody. Following labeling, antibodies were diluted in Candor PBS Antibody Stabilization solution (Candor Bioscience GmbH, Wangen, Germany) to 0.2 mg/ml and stored long term at 4 °C.

### Cell Culture

Glioblastoma U251-tGFP cells (a misidentified U251 line determined to be genetically identical to U373 by the ATCC; tGFP introduced by lentiviral transfection with multiplicity of 10, generously provided by S. Kumar’s Lab), were authenticated by short tandem repeat analysis and tested negative for mycoplasma. The U251-tGFP cells were maintained in a humidified 37 °C incubator kept at 5% CO_2_ with DMEM + Glutamax media (ThermoFisher 10566016) supplemented with 1× MEM nonessential amino acids (11140050, Life Technologies), 1% penicillin/streptomycin (15140122, Invitrogen), 1 mM sodium pyruvate (11360−070, Life Technologies), and 10% Fetal Bovine Serum (FBS, Gemini Bio-Products, 100-106). Cells were detached with 0.05% Trypsin-EDTA (ThermoFisher 25300-120) and resuspended in 4 °C 1x phosphate-buffered saline (PBS, Thermo Fisher Scientific 10010023) to generate cell suspensions used for scWB and scIEF.

### Single Cell Western Blots

The scWBs were performed as previously described^10^ with a few modifications. The microwell PA gel was created by chemically polymerizing with APS and TEMED an 8 %T, 3.3 %C, 3 mM BPMAC PA gel precursor solution on an SU-8 mold with microposts (32 μm diameter, ~75 μm height; 1 mm spacing along electrophoretic separation axis, 400 μm spacing between separation lanes) sandwiched to a silanized glass microscope slide. A U251-tGFP cell suspension (~500,000 cells/mL in 1× PBS, 4 °C) was introduced to the PA gel surface, cells were settled by gravity into the microwells (10 min), and excess cells were washed off the gel with PBS. Microwells were visually inspected under brightfield to ensure the majority of wells with cells had single-cell occupancy. Cells were lysed (30 s) within the wells in a 55 °C lysis/electrophoresis buffer (1x RIPA: 0.5% SDS, 0.25% sodium deoxycholate, 0.1% Triton X-100, 0.5x Tris-glycine, as previously reported^10^), and the proteins were electrophoresed into the gel at 40 V/cm (20s) in a custom electrophoresis chamber. Protein photo-immobilization was induced by application of UV at 100% intensity for 45 s with the Hamamatsu LC8 (Hamamatsu Photonics K.K.). Then, gels were rinsed in TBS-T for 30 min to remove uncaptured species.

### Single Cell Isoelectric Focusing

scIEF under denaturing conditions was performed as previously described^11^ with a few modifications. The microwell PA gel was created by chemically polymerizing with APS and TEMED a 6 %T, 3.3 %C, 3 mM BPMAC PA gel precursor solution on an SU-8 mold with microposts (32 μm diameter, ~40 μm height; single row of microwells positioned at a 2.25 mm distance from the acid region within the 9 mm focusing region, 500 μm spacing between separation lanes) sandwiched to a silanized glass microscope slide. Cell settling was performed as in the scWB. During cell settling, a three-component IEF lid gel was fabricated containing an acidic, focusing, and basic region. Supplemental Table T1 lists the components of the lid gel, which was polymerized for 4 min for each region at 20 mW/cm^2^ light intensity using a 390 nm UV long- pass filter (Edmund Optics) on an OAI model 30 collimated UV light source. The PA gel and lid gel were assembled in the ABS electrophoresis device as previously described^11^. After a 30 s delay for the lysis/focusing reagents in the focusing lid gel to diffuse into the microwell PA gel, IEF was conducted by applying 600 V for 6 min. Then, protein photo-immobilization and gel rinsing were performed as in the scWB.

### Immunoprobing

The gels were immunoprobed for tGFP as previously described^15^. Briefly, gels were exposed to 40 μL (scWB) or 12 μL (scIEF) of 33 μg/ml primary rabbit-anti-tGFP antibody in 2% BSA/TBS-T (metal-tagged or untagged, depending on the experiment) for 2 hrs, washed with TBS-T 2x for 30 min, exposed to 67 μg/ml secondary donkey-anti-rabbit-647 antibody in 2% BSA/TBS-T, and washed with TBS-T 2x for 30 min. The gels were then rinsed briefly in DI water to remove salts and dried with a nitrogen stream before imaging.

### Fluorescence Microarray Scanner Micrograph Acquisition

Gels were imaged on the GenePix 4300A microarray scanner (Molecular Devices) for expressed tGFP fluorescence with the 488-filter set and immunoprobed fluorescence signal with the 647-filter set.

### Confocal Micrograph Acquisition

We used confocal imaging to measure the tGFP protein depth distribution in scWB protein bands. After scWB, a no. 1.5H glass coverslip (Ibidi 0107999097) was placed on top of the hydrated PA gel and placed coverslip side down onto the microscope stage. Confocal imaging experiments were conducted on an inverted Zeiss LSM 710 AxioObserver at the CRL Molecular Imaging Center. Images were acquired at room temperature using a 40× water immersion objective (LD C-Apochromat 40×/1.1 NA W Corr M27, Zeiss). tGFP was imaged using a 488nm laser at 100% power, using the MBS488/561/ 633 beam splitter and the Zen 2010 software (Zeiss). We collected fluorescence image stacks (field of view: 212.55 μm × 212.55 μm; cubic voxels: 1.66 μm × 1.66 μm × 1.30 μm).

### MIBI-TOF Micrograph Acquisition

To increase sample conductivity, the scIEF slide was coated with 15 nm gold (99.999% purity) using a sputter coater. The custom built MIBI-TOF tissue analyzer was operated as previously described^9^.

### Profilometry

Gel height was assessed with a Veeco Dektak 8M Stylus Profilometer. The PA gel was dehydrated at the time of measurement.

### Image/Micrograph Analysis and Quantitation

scWB micrographs were analyzed using in-house ImageJ and Matlab (R2019b, MathWorks) scripts as previously described^15^. Area under the curve (A.U.C.) fluorescence was calculated by curve-fitting the scWB bands (both the detection antibody and expressed tGFP fluorescence bands) to a Gaussian function and summing the intensity values between four standard deviations of the peak center. A.U.C. was only reported for scWB bands with a Gaussian fit R-squared value >0.7, for accurate selection of peak boundaries. Statistical analysis was carried out with custom and existing Matlab functions. Analysis of confocal data is described in Note S1. Analysis of MIBI-TOF data to produce images of scIEF, including background subtraction and denoising, was performed as previously described^16^.

## RESULTS AND DISCUSSION

### Metal-tagged antibody probes are compatible with in-gel single-cell immunoassays

We first sought to investigate whether and to what extent metal-tagging affects antibody performance in in-gel immunoassays. For mass spectrometry imaging approaches (e.g., MIBI-TOF) the first step is to stain the sample with a panel of metal-tagged probes^17,18^. Therefore, to use MIBI-TOF as a detection method for single-cell electrophoretic assays such as scWB and scIEF, immunoprobing needs to be performed with metal-tagged primary antibodies instead of the conventional untagged primary and fluorophore-tagged secondary antibody probe duo. We hypothesized that metal-tagging may impact the physiochemical properties of an antibody molecule to the detriment of in-gel immunoassay performance. While previous studies have validated metal-tagged antibodies perform qualitatively similar to untagged antibody probes in fixed tissue^19,20^, the impact metal-tagging has on antibody probe performance in PA gel has not been studied.

To understand the performance of untagged versus metal-tagged anti-tGFP antibodies in the scWB assay, we performed indirect detection of the primary anti-tGFP antibody with a fluorophore-tagged secondary antibody. While fluorescence readout lacks the multiplexing capability of MIBI-TOF, we use fluorescence readout here as validation of immunoprobing of scIB with metal-tagged antibodies (Figure 1A). The basic steps of scIB are described in Figure 1B. We incubated scWB chips with either untagged rabbit anti-tGFP primary antibody or with Holmium-tagged rabbit anti-tGFP primary antibody (Figure 2A). Since the same polymer chemistry can be used for a large swath of metal isotopes, we employed the Holmium-tagged anti-tGFP antibody as the representative metal-tagged antibody. Both the untagged and metal-tagged anti-tGFP antibodies were selective for tGFP in the scWB as indicated by the overlapping protein bands shown in the micrographs and intensity plots (Figure 2B). We also tested the metal-tagged antibodies in scIEF, where denaturing conditions render the native tGFP signal undetectable, and the metal-tagged antibodies yielded qualitatively similar micrographs of the three tGFP proteoforms versus untagged antibodies (Figure 2C), which is aligned with previous work separating tGFP proteoforms with scIEF11. Notably, scWB does not resolve the three tGFP proteoforms that are observed with scIEF under denaturing conditions; therefore, tGFP appears as a single protein band in scWB.

**Figure 1.**
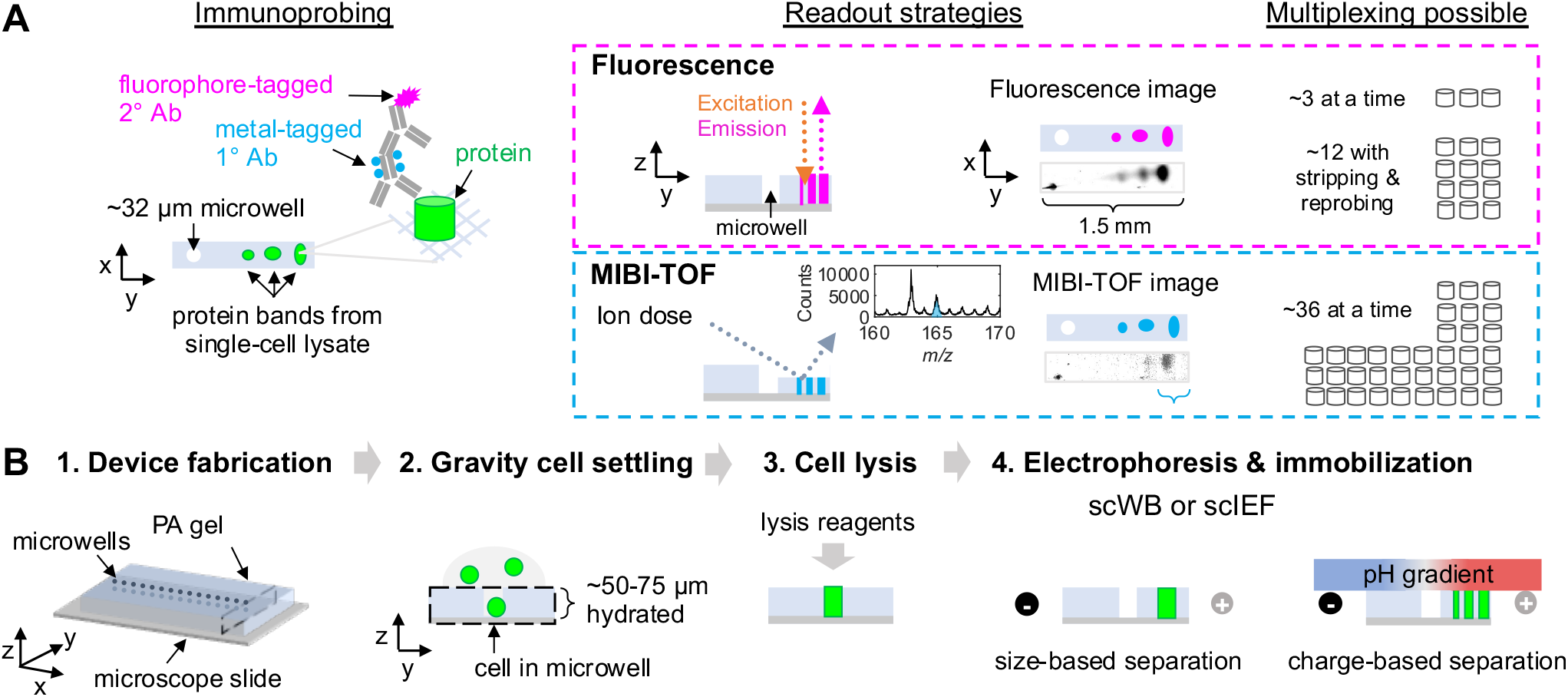
scIB-MIBI-TOF combines single-cell protein separation assays with multiplexed mass spectrometry detection. (A) During the immunoprobing step, the scIB sample is incubated with metal-tagged primary antibody (1° Ab), followed by fluorophore-tagged secondary antibody (2° Ab). The scIB sample is then imaged by both fluorescence and MIBI-TOF readout strategies, which differ in their multiplexing capability. In MIBI-TOF inset, mass spectrum is from summed counts from region indicated by the blue brace. (B) scIB is performed via the following steps: (1) microwell patterned PA gel is grafted to a microscope slide, (2) individual cells are settled on the hydrated PA gel matrix, (3) lysis reagents are introduced, and (4) an electric field is applied for electrophoresis and UV light is applied to activate a photoactive moiety in the PA gel backbone in order to covalently attach proteins to the PA gel (immobilization). A lid gel is introduced in scIEF to establish a pH gradient to separate proteins based on charge.

**Figure 2.**
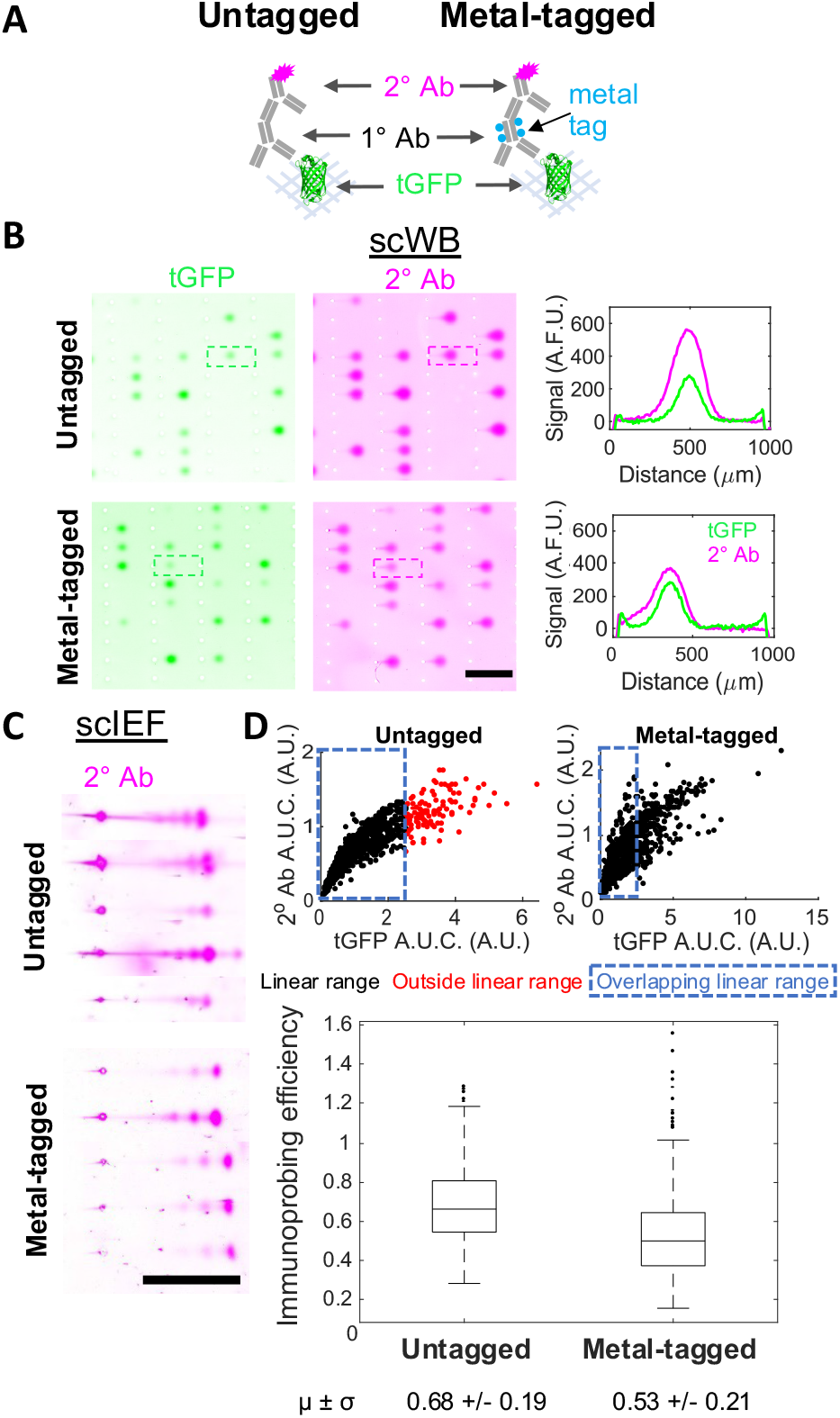
Metal-tagged antibody performance in scWB and scIEF. (A) Immunoprobing scheme. (B) Fluorescence images and intensity plots of scWBs of U251-tGFP cells probed for tGFP with untagged versus metal-tagged primary antibody (1° Ab), followed by a fluorophore-tagged secondary antibody (2° Ab). Both expressed tGFP and 2° Ab signal displayed. Scale bar is 1 mm. Micrographs in the same channel have the same acquisition settings, brightness, & contrast (representative micrographs from n_Untagged_ = 4, n_Metal-tagged_ = 4 independent scWB chips). (C) Fluorescence images of scIEF of U251-tGFP cells probed as in (B). Only 2° Ab signal displayed. Scale bar is 1 mm. Micrographs have the same acquisition settings, brightness, & contrast within each condition but not between the two conditions for better visualization (representative micrographs from n_Untagged_ = 2, n_Metal-tagged_ = 5 independent scIEF chips). (D) Scatter plots of 2° Ab A.U.C. versus tGFP A.U.C. with the linear data indicated in black, nonlinear data indicated in red, and dashed blue box surrounding overlapping linear data used to generate box plot of immunoprobing efficiency for untagged and metal-tagged configurations. Horizontal line in the box is the median (higher for gels immunoprobed with untagged 1° Ab, Mann−Whitney U-test p-value <0.0005) and box edges are at 25th and 75th percentile. Mean and standard deviation of data is displayed below plot. n_Untagged_ = 849 cells, n_Metal-tagged_ = 728 cells from 4 independent scWB chips for each condition.

We next characterized the relative immunoprobing efficiency of the untagged versus metal-tagged anti-tGFP antibody. Here we define immunoprobing efficiency as the ratio of probed A.U.C. to expressed tGFP A.U.C. We calculated immunoprobing efficiency for protein bands after determining the linear range by using an established approach^21^ to exclude high expression protein bands that were in an antibody-limited regime. We measured an immunoprobing efficiency that was 22% lower for the metal-tagged primary antibody configuration, as compared to the untagged primary antibody configuration (reduction from 0.68 in untagged to 0.53 for metal-tagged) (Figure 2D).

We attribute the slight reduction in immunoprobing efficiency for the metal-tagged antibody configuration to one or a combination of the following effects: (1) reduced primary-target binding efficiency, (2) reduced metal primary antibody partitioning into gel, or (3) reduced primary-secondary binding efficiency. Effect (3) is irrelevant to scIB-MIBI-TOF, as the secondary antibody is employed here to allow characterization even when an untagged primary antibody is of interest. Metals can be conjugated to IgG antibodies using either monomeric or polymeric bifunctional chelating agents (BFCAs) via sulfhydryl chemistry. Polymeric BFCAs (which were what was employed here using the MIBItag Conjugation Kit for metal-tagging) offer superior metal-tagging of antibodies because each repeating unit offers an opportunity to form a complex with a metal ion^22^. However, metal-tagging adds mass to the already bulky antibody probe. Added mass, and therefore a potential increase in hydrodynamic radius, would be expected to exacerbate antibody probe exclusion from the hydrogel (thermodynamic partitioning), thus further reducing the local, in-gel antibody concentration^23,24^ (effect (2)). Moreover, the metal-tag can interfere with the binding of antibody probe to antigen epitope^22^ (effect (1)).

Therefore, the 22% reduction represents a worst-case scenario for immunoprobing efficiency for this representative example (8 %T 3.3 %C PA gel, Holmium-tagging of an anti-tGFP antibody). For reference, in the stripping and reprobing multiplexing strategy, our group has previously reported a ~75% reduction in antibody signal after one round of stripping^13^. Additionally, the photoactive and hydrophobic moiety used to immobilize proteins in the PA gels after electrophoresis (BPMAC) has been shown to cause nonspecific retention of unbound antibody probes^24^, which we hypothesized could lead to increased background signal arising from any additional interaction between the metal-tag and the BPMAC. However, the metal-tagged antibody configuration did not increase background signal intensity (Figure S1). Altogether, these results indicate that metal-tagged antibody probes are compatible with detection of protein targets embedded in PA gel. Moreover, the indirect detection of metal-tagged primary antibodies with a fluorophore-tagged secondary antibody is a useful strategy to validate metal-tagged probes prior to incorporating the probes for MIBI-TOF detection.

### Protein signal detected increases with increasing depth imaged

To achieve MIBI-TOF readout of scIB assays, metal atoms from metal-labeled proteins embedded in a ~3.5-μm thick dehydrated PA gel must be ionized for down-stream mass spectrometry analysis, since MIBI-TOF is a SIMS instrument^9^. The basis of SIMS is the sputtering process in which the sample is ionized layer-by-layer beginning from the top of the sample to the bottom^25^. Primary ions from an ion source penetrate the sample surface, transferring energy to the sample through a collision cascade, which then causes secondary ions (mono- and polyatomic) to be ejected from the surface, exposing new surface^14^. Sample imaging requires sustained or repeated bombardment of the sample surface until the desired depth has been ionized and detected^26^. Consequently, MIBI-TOF images thin layers that can be used to reconstruct a final 3D image (analogous to confocal microscopy), whereas conventional fluorescence microarray scanners used to image scIB simultaneously integrate a wide depth of field to generate a single 2D image. Accordingly, for MIBI-TOF of scIB, the sample needs to be treated as a 3D substrate. To that end, we sought to characterize the depth distribution of protein signal in scIB assays to determine the depth at which we could expect to attain the maximum signal-to-noise (SNR) ratio; in other words, what percentage of sample to rasterize for optimal MIBI-TOF detection.

In the conventional MIBI-TOF imaging workflow, only a thin surface layer (~200 nm) of the 4-μm thick fixed tissue slice is typically imaged^9,27^. However, in the scIB system, diffusion during lysis and electrophoresis may dilute protein in scIB samples more than in fixed tissue, so we hypothesized imaging of scIBs would require deeper sample imaging. Though lateral diffusion of protein signal in scIBs has been well characterized^28^, the depth distribution of signal in scIB protein bands has only been computationally interrogated^29^. During electrophoresis, the motion of charged molecules in the direction of the electric field is governed by the electrostatic Coulomb force and opposite viscous drag^30^. However, the driving force in the z-direction is the concentration gradient, which causes protein loss as proteins diffuse and partition between the PA gel and the fluid and/or gel lid above the device (Figure 3A schematics). Based on diffusional loss of protein out of the microwell during cell lysis and out of the PA gel during electrophoresis, we hypothesized that the protein signal will be concentrated towards the PA gel-microscope slide interface (“bottom of the gel”).

**Figure 3.**
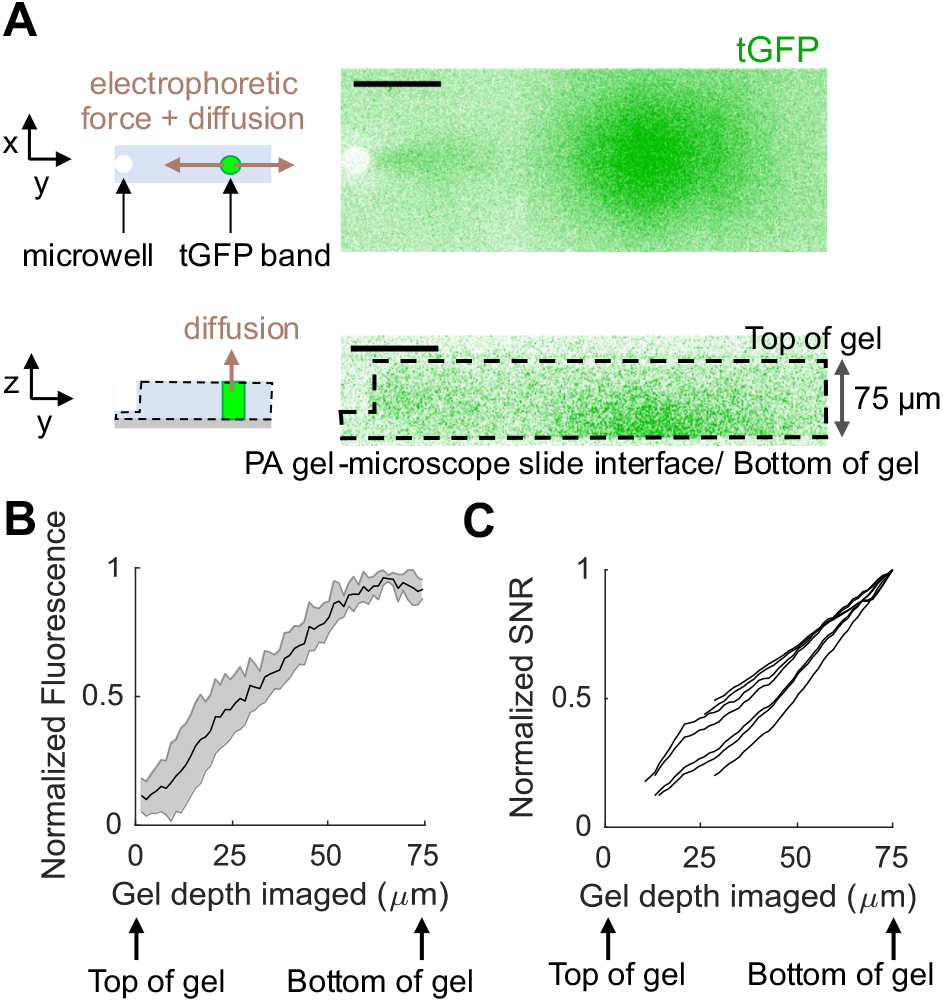
Protein signal is concentrated towards the bottom of the PA gel in scIB. (A) Schematics depict the forces acting on protein molecules during electrophoresis. Micrographs are a top (x-y) and side (z-y) view of a single scWB separation lane of U251-tGFP cells imaged with fluorescence confocal microscopy. tGFP is in green. Scale bar is 100 μm. (B) The plot is the normalized fluorescence intensity after background subtraction of tGFP bands from confocal z-stack images over the gel depth, shaded error region is standard deviation. (C) Each line in this plot represents the normalized SNR for a single cell as a function of how much percentage of the gel was included in the SNR measurement (Note S1). For (B) and (C), n = 7 cells from 2 independent scWB chips. Cell-to-cell variation resulted in large differences in absolute fluorescence intensity and SNR values, so normalization to the maximum fluorescence intensity and SNR value, respectively, within each cell allowed improved side-by-side comparison of the biological replicates.

To experimentally determine the depth protein concentration in a scIB assay, we directly imaged tGFP bands in a hydrated scWB chip with fluorescence confocal microscopy. Figure 3A shows a top view of a tGFP band and the corresponding side view showing the underlying depth protein distribution. As we hypothesized, the tGFP signal is concentrated at the bottom of the gel with protein concentration going to zero at the top of the gel (Figure 3B).

We next sought to understand how the depth protein distribution in scIB would impact SNR in MIBI-TOF. To approximate the MIBI-TOF process of sputtering beginning at the top of the gel and sputtering increasing layers, we added increasing numbers of z-stack fluorescence confocal slices and calculated the SNR of the protein band in each summed image (Note S1). By excluding images that yielded SNR < 3 and plotting SNR normalized to the maximum SNR from the series of summed images, we see that the first few layers of the gel are insufficient to yield an SNR greater than 3 and the entirety of gel depth (~75 μm) should be imaged to reach the maximum SNR (Figure 3C). Since SNR is directly proportional to gel depth imaged, we can anticipate that, to improve the detection of low SNR (low abundance) protein targets with MIBI-TOF (as opposed to the fluorescence confocal microscopy used here), the depth of gel imaged should be as close to the total gel height as possible.

### Modulating gel depth rasterized by changing MIBI-TOF ion dose

We next characterized the relationship between depth of PA gel rasterized and ion dose. Ion dose is a function of imaging parameters that can be adjusted in the MIBI-TOF instrument (equation 1, *Ion dose* = area normalized ion dose, *I*=primary ion current, *t*=acquisition time for a single depth, *d*=depths acquired, *A*=field area in mm^2^).

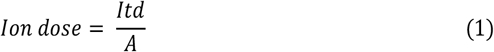

Since SIMS is performed in a vacuum chamber, samples for MIBI-TOF are dehydrated before insertion into the instrument. Dehydrated scIB gels (~3.5 μm) have a similar thickness to the tissue sections (4 μm) employed in previous MIBI-TOF studies^9,16^. The entire depth of a 4-μm thick tissue section has been previously imaged with MIBI-TOF^9^. To access the proteins that would be embedded in the 3.5-μm thick scIB chip, we sought to determine the ion dose required to ionize and image various PA gel depths.

Figure 4A shows a profilometer trace of MIBI-TOF imaged spots on 6 %T PA gel at various ion doses. The depth rasterized was measured using a stylus profilometer which physically drags a stylus across the gel surface to generate a trace of the depth profile. The highest ion dose tested rasterized ~50% of the gel depth with an ion dose of 80 nA×hr/mm^2^. We observed a linear relationship between depth rasterized and ion dose (Figure 4B, R^2^ = 0.8226). The sputter yield of individual species increases linearly with applied ion flux^14^, so the linear relationship between depth rasterized and ion dose suggests that all species in the PA gel sample are being sputtered at nearly the same rate. Future work will determine whether this constant erosion rate is maintained when imaging the entire 3.5 μm PA gel. Notably, a tradeoff between imaging throughput and detection sensitivity is expected, because the higher ion dose images (that rasterize deeper into the gel) have the potential to increase SNR (Figure 3C) yet require longer acquisition times (equation 1).

**Figure 4.**
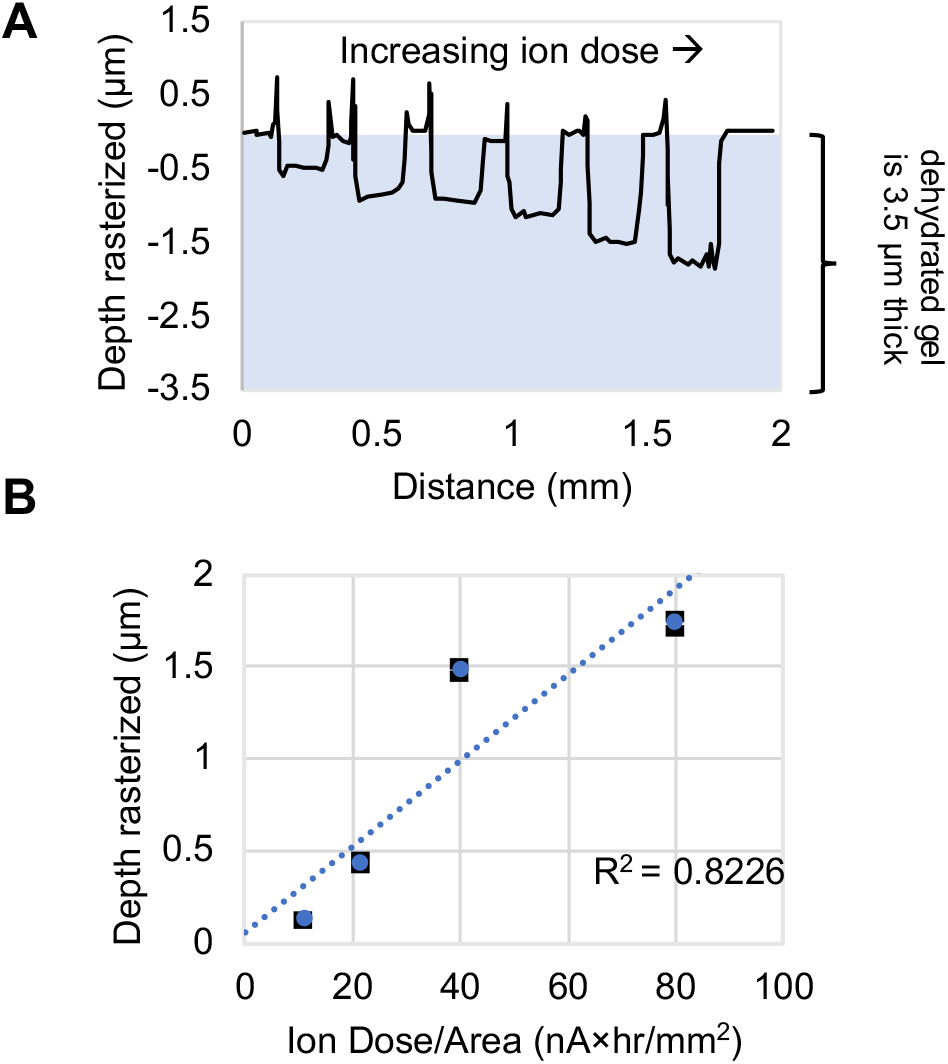
PA gel depth rasterized can be tuned by modulating ion dose. (A) Profilometer trace of MIBI-TOF imaged spots with increasing ion dose from left to right. (B) Plot of depth rasterized vs. ion dose applied (black error bards are the standard deviation and may be smaller than blue data point symbols, n >= 4 imaged spots per ion dose on same PA gel). See Table S2 for imaging conditions.

### MIBI-TOF of scIEF resolves tGFP proteoforms from single cells

To validate MIBI-TOF for scIB, we compared MIBI-TOF images to fluorescence images using the same immunoprobing scheme used in Figure 2A for the metal-tagged configuration. Figure 5A shows the scIEF images of tGFP proteoforms *α*, *β*, and *γ*, as detected with the fluorescence and with MIBI-TOF, respectively. The corresponding intensity profiles are shown. As expected, we observed correlation between fluorescence and MIBI-TOF readouts (Figure 5B with colocalized signal in black), but MIBI-TOF was unable to detect the lowest abundance proteoform, *γ*. At the ion dose used for this acquisition, ~42% of the gel was rasterized, which suggests there may be insufficient metal-tagged antibody probe in the depth ionized in this acquisition to produce detectable signal for proteoform *γ*. Even at the lowest resolution settings possible on this MIBI-TOF instrument, the instrument was set up for nanometer-scale tissue analysis. The application here only requires a resolution of 10s of micrometers. As such, we expect a lower resolution instrument configuration would have exponentially higher primary ion beam power, thus be able to sample more gel for better SNR over a shorter period of time (acquisition time of the micrograph for Well 1 in Figure 5A was ~35 minutes). Moreover, there remains avenues of sample preparation optimization to increase substrate conductivity (in addition to or instead of the 15 nm gold coating), and thus, sensitivity of detection of the secondary reporter ions. Altogether, these results demonstrate successful MIBI-TOF detection of two distinct tGFP proteoforms separated using scIEF, a scIB assay.

**Figure 5.**
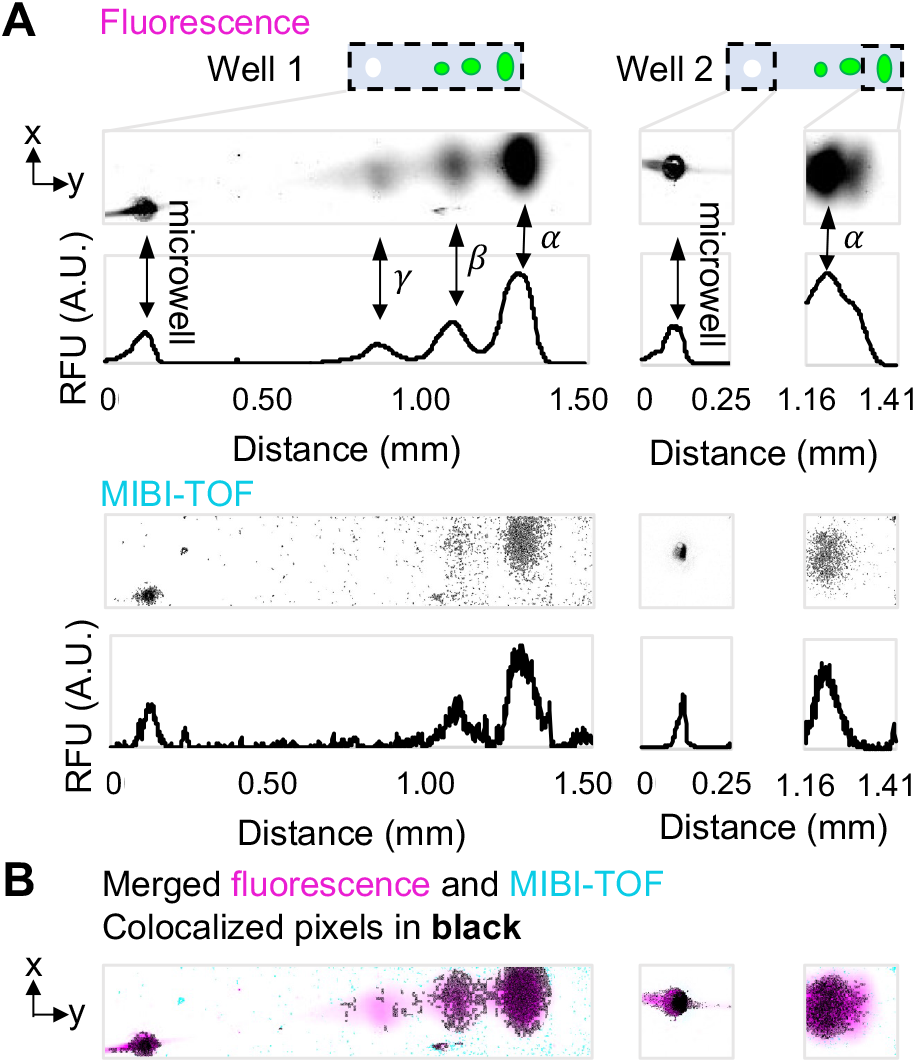
MIBI-TOF of scIEF resolves tGFP proteoforms from single cells. (A) Fluorescence vs. MIBI-TOF micrographs and intensity plots of same separated tGFP proteoforms (proteoforms are denoted α, β, and γ) from U251-tGFP cells. Well 1 MIBI-TOF image is composed of 8 tiled images of ~1 single cell separation. Well 2 MIBI-TOF image is composed of 2 tiled images of ~1 single cell separation. (B) Colocalized pixel map of merged images. See Table S2 for imaging conditions. The x-axis of intensity plots is also the scale bar for micrographs in (B) and (C).

## CONCLUSIONS

The MIBI-TOF-based single-cell immunoblotting performance reported here forms a promising basis for the extension of MIBI-TOF readouts to other bioanalytical assays and samples where multiplexed detection from a 3D matrix is desirable. In the case of single-cell immunoblotting, by demonstrating the feasibility of MIBI-TOF readout, we increased the amount of simultaneously available antibody labels from ~3 to ~40 and eliminated the need to perform antibody stripping and reprobing for multiplexed target detection, including for proteoforms. scIB-MIBI-TOF provides a promising strategy to increase the number of low-abundance targets detected with a simplified experimental workflow. Importantly, due to the spatial separation between protein bands, we detected two distinct tGFP proteoforms with MIBI-TOF, yet only one metal-tag channel was utilized. Besides the ~40 channels provided by the distinct metal-tags in MIBI-TOF, scIB provides an additional opportunity to increase the current multiplexing capability of MIBI-TOF by a factor of approximately the peak capacity of the scIB assay. Peak capacity is the number of theoretical protein bands that can “fit” in a separation lane, which is ~10 for scWB with a 1 mm separation lane^31^ and ~17 for scIEF with a 9 mm separation lane^11^. Building on these results, ongoing research is focusing on multiplexed detection in additional channels by utilizing additional metal-tagged antibodies during immunoprobing.

## ASSOCIATED CONTENT

### Supporting Information

The Supporting Information is available free of charge on the ACS Publications website.

Background signal for metal-tagged antibodies, Calculation of normalized SNR, Composition of IEF lid gel, Imaging conditions and depth rasterized data (PDF)

## AUTHOR INFORMATION

### Author Contributions

All authors designed the experiments. G.L. and M.B. performed the experiments. G.L. performed the data analysis. All authors wrote the manuscript. All authors have given approval to the final version of the manuscript.

### Notes

The authors declare the following competing interest(s): Co-authors may benefit financially from licensing or royalties stemming from University-owned intellectual property on the underlying technologies. M.A. and S.C.B. are consultants and shareholders in IonPath Inc.

## ACKNOWLEDGMENT

This work was funded in part by the National Institutes of Health grant R01CA203018, National Cancer Institute of the National Institutes of Health, Cancer Moonshot award, Grant Number: 1R33CA225296-01, and by the Chan Zuckerberg Biohub to A.E.H. M.A. was supported by 1-DP5-OD019822. S.C.B. and M.A. were jointly supported by 1R01AG056287 and 1R01AG057915, 1U24CA224309, and the Bill and Melinda Gates Foundation. G.L. gratefully acknowledges support from the National Science Foundation Graduate Research Fellowship Program (NSF GRFP). Photolithography and profilometry was performed in the QB3 Biomolecular Nanotechnology Center. Confocal imaging experiments were conducted at the CRL Molecular Imaging Center at UC Berkeley, supported by the Gordon and Betty Moore Foundation. We acknowledge all members of the Herr Lab at UC Berkeley for useful discussions and feedback, and especially Heather Robison, Ph.D. for initiating the collaboration and Alisha Geldert for advice on analysis.

## REFERENCES

(1) Yang, L.; George, J.; Wang, J. Deep Profiling of Cellular Heterogeneity by Emerging Single-Cell Proteomic Technologies. PROTEOMICS 2020, 20 (13), 1900226. https://doi.org/10.1002/pmic.201900226.

(2) Aebersold, R. et al. How Many Human Proteoforms Are There? Nat. Chem. Biol. 2018, 14 (3), 206–214. https://doi.org/10.1038/nchembio.2576.

(3) Smith, L. M.; Kelleher, N. L. Proteoform: A Single Term Describing Protein Complexity. Nat. Methods 2013, 10 (3), 186–187. https://doi.org/10.1038/nmeth.2369.

(4) Arribas, J.; Baselga, J.; Pedersen, K.; Parra-Palau, J. L. P95HER2 and Breast Cancer. Cancer Res. 2011, 71 (5), 1515–1519. https://doi.org/10.1158/0008-5472.CAN-10-3795.

(5) Levy, E.; Slavov, N. Single Cell Protein Analysis for Systems Biology. Essays Biochem. 2018, 62 (4), 595–605. https://doi.org/10.1042/EBC20180014.

(6) Chattopadhyay, P. K.; Roederer, M. Cytometry: Today’s Technology and Tomorrow’s Horizons. Methods 2012, 57 (3), 251–258. https://doi.org/10.1016/j.ymeth.2012.02.009.

(7) Regnier, F. E.; Kim, J. Proteins and Proteoforms: New Separation Challenges. Anal. Chem. 2018, 90 (1), 361–373. https://doi.org/10.1021/acs.analchem.7b05007.

(8) Dickinson, M. e.; Bearman, G.; Tille, S.; Lansford, R.; Fraser, S. e. Multi-Spectral Imaging and Linear Unmixing Add a Whole New Dimension to Laser Scanning Fluorescence Microscopy. BioTechniques 2001, 31 (6), 1272–1278. https://doi.org/10.2144/01316bt01.

(9) Keren, L.; Bosse, M.; Thompson, S.; Risom, T.; Vijayaragavan, K.; McCaffrey, E.; Marquez, D.; Angoshtari, R.; Greenwald, N. F.; Fienberg, H.; Wang, J.; Kambham, N.; Kirkwood, D.; Nolan, G.; Montine, T. J.; Galli, S. J.; West, R.; Bendall, S. C.; Angelo, M. MIBI-TOF: A Multiplexed Imaging Platform Relates Cellular Phenotypes and Tissue Structure. Sci. Adv. 2019, 5 (10), eaax5851. https://doi.org/10.1126/sciadv.aax5851.

(10) Hughes, A. J.; Spelke, D. P.; Xu, Z.; Kang, C.-C.; Schaffer, D. V.; Herr, A. E. Single-Cell Western Blotting. Nat. Methods 2014, 11 (7), 749–755. https://doi.org/10.1038/nmeth.2992.

(11) Tentori, A. M.; Yamauchi, K. A.; Herr, A. E. Detection of Isoforms Differing by a Single Charge Unit in Individual Cells. Angew. Chem. Int. Ed. 2016, 55 (40), 12431–12435. https://doi.org/10.1002/anie.201606039.

(12) Sinkala, E.; Sollier-Christen, E.; Renier, C.; Rosàs-Canyelles, E.; Che, J.; Heirich, K.; Duncombe, T. A.; Vlassakis, J.; Yamauchi, K. A.; Huang, H.; Jeffrey, S. S.; Herr, A. E. Profiling Protein Expression in Circulating Tumour Cells Using Microfluidic Western Blotting. Nat. Commun. 2017, 8, 14622. https://doi.org/10.1038/ncomms14622.

(13) Gopal, A.; Herr, A. E. Multiplexed In-Gel Microfluidic Immunoassays: Characterizing Protein Target Loss during Reprobing of Benzophenone-Modified Hydrogels. Sci. Rep. 2019, 9 (1), 15389. https://doi.org/10.1038/s41598-019-51849-8.

(14) Vickerman, J. C. ToF-SIMS—An Overview. In TOF-SIMS: Surface Analysis by Mass Spectrometry; Manchester and IM Publications: Chichester, 2001; pp 1–40.

(15) Kang, C.-C.; Yamauchi, K. A.; Vlassakis, J.; Sinkala, E.; Duncombe, T. A.; Herr, A. E. Single Cell–Resolution Western Blotting. Nat. Protoc. 2016, 11. https://doi.org/10.1038/nprot.2016.089.

(16) Keren, L.; Bosse, M.; Marquez, D.; Angoshtari, R.; Jain, S.; Varma, S.; Yang, S.-R.; Kurian, A.; Van Valen, D.; West, R.; Bendall, S. C.; Angelo, M. A Structured Tumor-Immune Microenvironment in Triple Negative Breast Cancer Revealed by Multiplexed Ion Beam Imaging. Cell 2018, 174 (6), 1373–1387.e19. https://doi.org/10.1016/j.cell.2018.08.039.

(17) Baharlou, H.; Canete, N. P.; Cunningham, A. L.; Harman, A. N.; Patrick, E. Mass Cytometry Imaging for the Study of Human Diseases—Applications and Data Analysis Strategies. Front. Immunol. 2019, 10. https://doi.org/10.3389/fimmu.2019.02657.

(18) Yu, Y.; Dang, J.; Liu, X.; Wang, L.; Li, S.; Zhang, T.; Ding, X. Metal-Labeled Aptamers as Novel Nanoprobes for Imaging Mass Cytometry Analysis. Anal. Chem. 2020, 92 (9), 6312–6320. https://doi.org/10.1021/acs.analchem.9b05159.

(19) Angelo, M.; Bendall, S. C.; Finck, R.; Hale, M. B.; Hitzman, C.; Borowsky, A. D.; Levenson, R. M.; Lowe, J. B.; Liu, S. D.; Zhao, S.; Natkunam, Y.; Nolan, G. P. Multiplexed Ion Beam Imaging of Human Breast Tumors. Nat. Med. 2014, 20 (4), 436–442. https://doi.org/10.1038/nm.3488.

(20) Rost, S.; Giltnane, J.; Bordeaux, J. M.; Hitzman, C.; Koeppen, H.; Liu, S. D. Multiplexed Ion Beam Imaging Analysis for Quantitation of Protein Expresssion in Cancer Tissue Sections. Lab. Invest. 2017, 97 (8), 992–1003. https://doi.org/10.1038/labinvest.2017.50.

(21) Kroll, M. H.; Emancipator, K.; Floering, D.; Tholen, D. An Algorithm for Finding the Linear Region in a Nonlinear Data Set. Comput. Biol. Med. 1999, 29 (5), 289–301. https://doi.org/10.1016/S0010-4825(99)00011-6.

(22) Han, G.; Spitzer, M. H.; Bendall, S. C.; Fantl, W. J.; Nolan, G. P. Metal-Isotope-Tagged Monoclonal Antibodies for High-Dimensional Mass Cytometry. Nat. Protoc. 2018, 13 (10), 2121–2148. https://doi.org/10.1038/s41596-018-0016-7.

(23) Gehrke, S. H.; Fisher, J. P.; Palasis, M.; Lund, M. E. Factors Determining Hydrogel Permeability. Ann. N. Y. Acad. Sci. 1997, 831 (1), 179–184. https://doi.org/10.1111/j.1749-6632.1997.tb52194.x.

(24) Su, A.; Smith, B. E.; Herr, A. E. In Situ Measurement of Thermodynamic Partitioning in Open Hydrogels. Anal. Chem. 2020, 92 (1), 875–883. https://doi.org/10.1021/acs.analchem.9b03582.

(25) Nuñez, J.; Renslow, R.; Cliff, J. B.; Anderton, C. R. NanoSIMS for Biological Applications: Current Practices and Analyses. Biointerphases 2017, 13 (3), 03B301. https://doi.org/10.1116/1.4993628.

(26) Walker, A. V. Secondary Ion Mass Spectrometry. In Encyclopedia of Spectroscopy and Spectrometry; Elsevier, 2016; pp 44–49. https://doi.org/10.1016/B978-0-12-803224-4.00022-4.

(27) Levenson, R. M.; Borowsky, A. D.; Angelo, M. Immunohistochemistry and Mass Spectrometry for Highly Multiplexed Cellular Molecular Imaging. Lab. Invest. 2015, 95 (4), 397–405. https://doi.org/10.1038/labinvest.2015.2.

(28) Tan, K. Y.; Herr, A. E. Ferguson Analysis of Protein Electromigration during Single-Cell Electrophoresis in an Open Microfluidic Device. Analyst 2020, 145 (10), 3732–3741. https://doi.org/10.1039/C9AN02553G.

(29) Rosàs-Canyelles, E.; Dai, T.; Li, S.; E. Herr, A. Mouse-to-Mouse Variation in Maturation Heterogeneity of Smooth Muscle Cells. Lab. Chip 2018, 18 (13), 1875–1883. https://doi.org/10.1039/C8LC00216A.

(30) Kirby, B. J. Micro- and Nanoscale Fluid Mechanics: Transport in Microfluidic Devices; Cambridge University Press, 2010.

(31) Yamauchi, K. A.; Herr, A. E. Subcellular Western Blotting of Single Cells. Microsyst. Nanoeng. 2017, 3 (1), 1–9. https://doi.org/10.1038/micronano.2016.79.

