## Supplementary Information for "Multiplexed Ion Beam Imaging Readout of Single-Cell Immunoblotting"

^1^The UC Berkeley/UCSF Joint Graduate Program in Bioengineering

^2^Department of Bioengineering, University of California, Berkeley, California 94720, United States ^3^Department of Pathology, Stanford University, Stanford, California 94025, United States

**Table of Contents**

**Supplemental Figure S1. Background signal for metal-tagged antibodies…………………………...S-2**

**Supplemental Note S1. Calculation of normalized SNR…………………………………………………..S-3**

**Supplemental Table S1. Composition of IEF lid gel……………………………………………………….S-4**

**Supplemental Table S2. Imaging conditions and depth rasterized data………………………………S-5**

**Reference………………………………………………………………………………………………………….S-6**

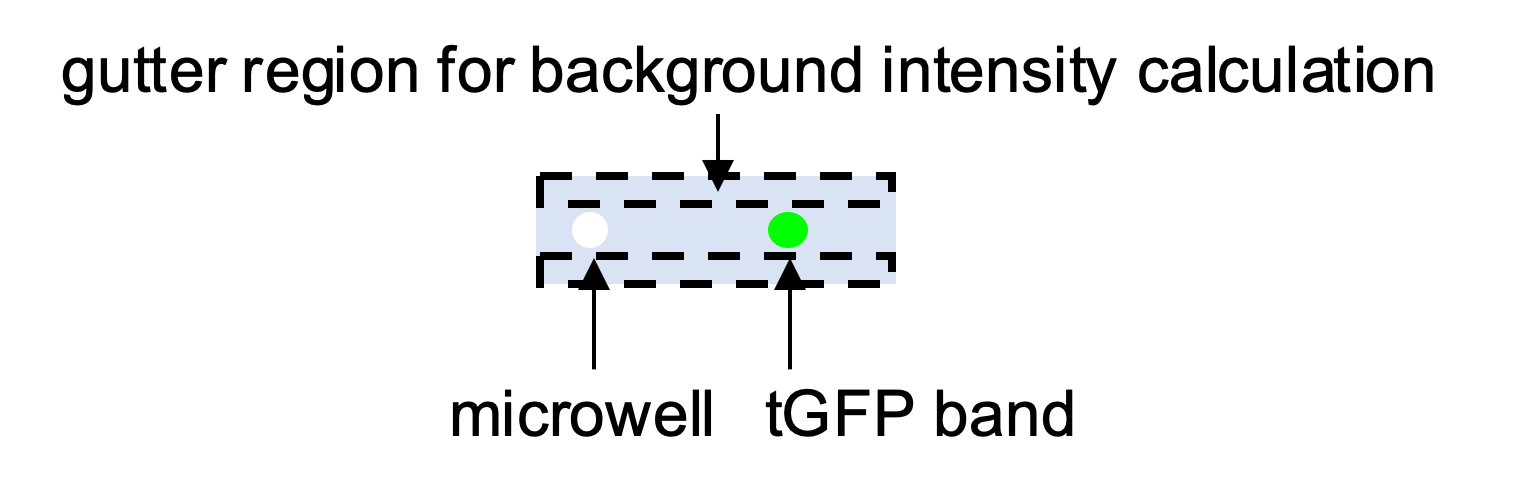

untagged

metal-tagged

0

0.2

0.4

0.6

0.8

1

1.2

Background (A.F.U.)

| µ ± σ | 0.161 +/- 0.062 | 0.156 +/- 0.064 |
| --- | --- | --- |

**Supplemental Figure S1. Background signal for metal-tagged antibodies.** Boxplot of background signal from the same results depicted in main text Figure 2D. Background intensity was calculated from a gutter region of each analyzed ROI chosen as 3-4 standard deviations away from the peak center as depicted in the schematic. Horizontal line in the box is the median (higher for gels immunoprobed with untagged 1° Ab, Mann−Whitney U-test p-value <0.0005) and box edges are at 25th and 75th percentile. Mean and standard deviation of data is displayed below plot. Difference in background signal in scWB for metal-tagged antibody versus untagged antibody configuration is statistically significant, but small. n_Untagged_ = 849 cells, n_Metal-tagged_ = 728 cells.

**Note S1. Calculation of normalized SNR**

We calculated a normalized SNR in Figure 3C of increasingly summed confocal slices to investigate the relationship between percent of gel depth imaged and SNR, since percent of gel depth imaged is a tunable parameter in MIBI-TOF. With an in-house Matlab script, we performed the following:

1. From each confocal z-stack of a single scIB lane with n slices, we summed slices 1, 1-2, 1-3, 1-4…1-n with slice 1 being the top layer of the gel and slice n being the bottom layer of the gel (gel-microscope slide interface). The result was n images with the first image being just the top layer of the gel and the nth image being the sum of the entirety of the gel over its depth.
2. For each of the n images, we performed background subtraction, Gaussian fitting, and calculated SNR as previously described^1^. Images with protein bands with SNR < 3 were disregarded.
3. Since cell-to-cell variation resulted in large differences in absolute SNR values, we normalized each SNR value by the maximum SNR within the n images for each cell, which allowed improved side-by-side comparison of the biological replicates.

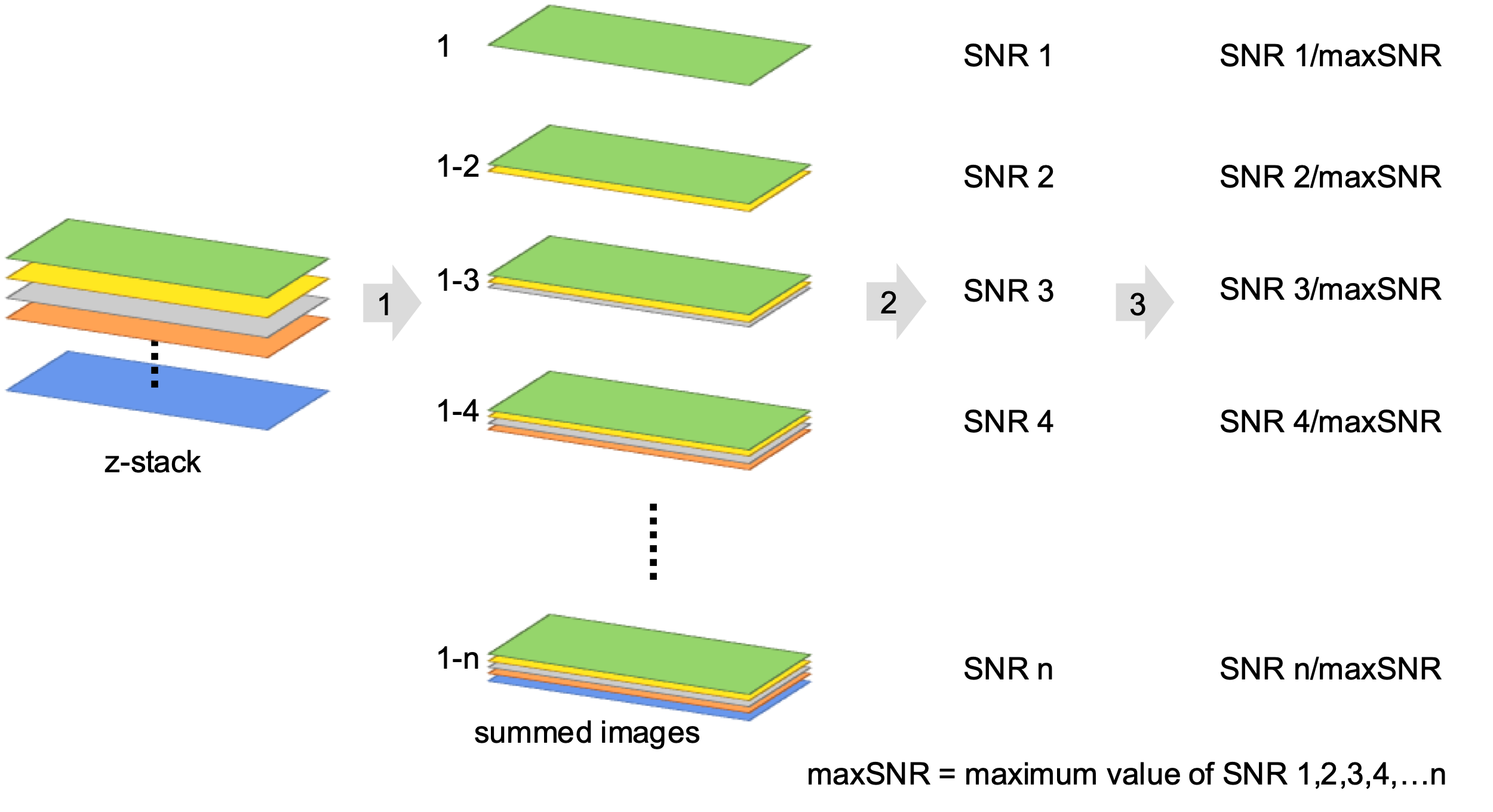
**Supplemental Table S1. Composition of IEF lid gel.** Components of the 3-part lid gel used for lysis and electrophoresis in the scIEF assay.

| **Lid gel components** | **pH 4 anolyte boundary condition** | **Focusing region** | **pH 10 catholyte boundary condition** |
| --- | --- | --- | --- |
| Polyacrylamide gel | - 15 %T - 3.3 %C - 0.2% VA-086 | - 15 %T - 3.3 %C - 0.2% VA-086 | - 15 %T - 3.3 %C - 0.2% VA-086 |
| IEF reagents and detergents |  | - 1% final ZOOM™ Carrier Ampholytes pH 4-7 - 1% (v/v) TritonX-100 - 3.6% (w/v) CHAPS - 0.0125% (w/v) digitonin - 7 M urea - 2 M thiourea |  |
| Boundary conditions | - 13.6 mM pKa 3.6 immobiline - 6.4 mM pKa 9.3 immobiline |  | - 5.6 mM pKa 3.6 immobiline - 14.4 mM pKa 9.3 immobiline |

| **Fig** | **Current (nA)** | **FOV size (µm)** | **dwell time (ms)** | **# planes** | **pixels** | **Ion dose/plane (nA**×**hr/mm^2^)** | **Ion dose total (nA**×**hr/mm^2^)** | **Average depth rasterized (µm)** |
| --- | --- | --- | --- | --- | --- | --- | --- | --- |
| **4B, 5** | 21.9 | 200 | 4 | 1 | 256 | 39.87 | 39.87 | 1.48 |
| **4B** | 21.9 | 200 | 4 | 2 | 256 | 39.87 | 79.74 | 1.73 |
| **4B** | 9.5 | 400 | 1 | 10 | 256 | 1.08 | 10.81 | 0.13 |
| **4B** | 47 | 400 | 4 | 1 | 256 | 21.39 | 21.39 | 0.44 |

**Supplemental Table S2. Imaging conditions and depth rasterized data.**

**Reference**

1.	Kang, C.-C. *et al.* Single cell–resolution western blotting. *Nat. Protoc.* **11**, (2016).
